# Cancer radiomic feature variations due to reconstruction kernel choice and integral tube current

**DOI:** 10.1101/2024.06.04.596806

**Authors:** Elfried Salanon, Anqi Fu, Aditya P. Apte, Usman Mahmoud, Zehor Belkhatir, Amita Shukla-Dave, Joseph O. Deasy

## Abstract

**Purpose:** Radiological cancer imaging features, or radiomics features, can be derived to diagnose disease or predict treatment response. However, variability between vendors, scanners, protocols, and even reconstruction software versions is an obstacle to the clinical use of radiomics features. This study aimed to characterize the impact of kernel reconstruction differences and integral tube current settings on radiomic features extracted from computed tomography (CT) scans.

**Methods:** Radiomic features were extracted from CT scans of a 3D-printed phantom with five imprinted tumors using the CERR software system, resulting in 282 features. Batch effects were assessed via principal component analysis (PCA) and correlation measures. Robustness was measured using the concordance correlation coefficient (CCC) and Pearson correlation coefficient. Statistical analysis was performed using R software.

**Results:** PCA identified two clusters comprised of Standard, ASIRs, ASIRV, and soft kernels in one, and Lung and Bone Kernels in the other. Features displayed a gradient from ASIR10 to ASIR50 and ASIRV1 to ASIRV5 in terms of nearness to Standard Kernel feature values. Feature correlation matrices revealed little change in ASIRs, ASIRVs, and the Standard Kernel, but showed significant changes in Bone and Lung Kernel results. Combat-algorithm correction improved robustness, particularly in first-order statistic features, and mitigated batch effects due to the ASIRs and the standard kernel. Forty (40) out of 282 features were identified as robust. However, Combat-based correction performed poorly in harmonizing Bone and Lung reconstruction kernels.

**Conclusions:** The robustness of means and median radiomic features across kernel reconstruction choices, in contrast to the lack of robustness in many other radiomic features, suggests that kernel reconstruction effects are not well-addressed by current harmonization methods.

## Introduction

High-throughput technologies have enabled the study of multiple biological pathways simultaneously, ushering in the era of “big data” in medical research. This is led by ‘omics’ fields like genomics, epigenomics, transcriptomics, proteomics, metabolomics and radiomics, which aim to achieve a better understanding of pathophysiology and an improvement in treatment and therapies for a variety of illnesses (Perng & Aslibekyan, 2020). However, one of the key challenges in ‘omics’ is repeatability and reproducibility. Difficulties in reproducing omics studies limit the generalizability of results and the translation of identified biomarkers and signatures to other settings (Goh et al., 2017; Goh & Wong, 2017; Perng & Aslibekyan, 2020). In some omics sciences, sample or cohort data is acquired in groups subject to identical or near identical conditions, referred to as batches. Oftentimes, differences are observed between batches, even when the same replicate is acquired. If deemed not due to sample variation, these differences are called batch effects and have been well-documented in genomics, proteomics, and metabolomics ((Liu et al., 2020; Phua et al., 2022; Tom et al., 2017). They affect statistical power and lead to false-positive results (Nygaard et al., 2016). Moreover, in an era of deep learning and artificial intelligence, batch effects have been observed to produce strong biases even when using cross-validation (Goh et al., 2017).

Radiomics is the extraction of features from radiographic imaging to quantify subtle changes in tissue structure and heterogeneity, which are imperceptible by visual assessment alone (Mayerhoefer et al., 2020). This approach harnesses the information in images to enhance a comprehensive characterization of tumor phenotypes (Aerts et al., 2014; Lambin et al., 2012). Radiomics studies have also been found to be impacted by different types of batch effects that occur at different steps of the feature extraction process (Castaldo et al., 2023; Cavinato et al., s. d.; Fornacon-Wood et al., 2020; Hertel et al., 2023; Ligero et al., 2021a; Paquier et al., 2022; van Velden et al., 2016; Zhao et al., 2014). Due to the multiple steps involved in the process, as well as a number of possibly changing factors *(equipment vendor, integral tube current, reconstruction kernels, slice thickness, etc.)*, and the fact that analyses are generally based on retrospective data, the definition, identification, and correction of batch effects in radiomics is particularly complex (Eshaghzadeh Torbati et al., 2021; Li et al., 2021; Tafuri et al., 2022).

Multiple studies have addressed this problem, and they can be classified into three categories. Some studies focus on the identification of sources of batch effects, such as the equipment vendor, reconstruction kernel, slice thickness, etc. (Haarburger et al., 2020; Ligero et al., 2021b, 2021a; Reiazi et al., 2021; Zhao et al., 2014). Other studies focus on the identification of radiomics features that are robust to batch effects, so they can be included in the radiomics signatures using the concordance and the correlations coefficient or the intraclass correlations coefficient (Carles et al., 2021; Lu et al., 2021). Finally, a body of work focuses on the development of batch effects correction methods and strategies for data harmonization and the minimization of acquisition-related variability (Cavinato et al., 2023; Eshaghzadeh Torbati et al., 2021; Horng et al., 2022; Lee et al., 2022; Ligero et al., 2021b; Mahon et al., 2020; Orlhac et al., 2022; Soliman et al., 2022). Considering the breadth of batch effects that have been identified and correction methods that have been proposed, it is clear that there is still a lack of consensus in the research community on the answers to several key questions:

1. *Given that multiple classes of features exist in radiomics, are all feature classes affected in the same way by batch effects?*
2. *If not, should we apply only one correction method to account for all the batch effects in radiomics or combine several?*
3. *How do corrections for different types of batch effects affect different classes of radiomics features?*

Addressing these questions is crucial for overcoming challenges related to batch effect correction and harmonization in radiomics. These barriers hinder the large-scale application and generalization of radiomics signatures. By finding solutions, we can develop better harmonization tools and improve data acquisition and analysis workflows. In this paper, we consider a batch to be a group of images sharing a set of acquisition parameters, and we consider a batch effect to be the non-biological/non-sample variation resulting from changes in the information production conditions. We evaluate the impact of two types of batch effects, namely the kernel batch effect and the integral tube current batch effect, on different classes of radiomics features and identify the radiomics features that are robust to these specific batch effects. By comparing the robust and non-robust features, we can characterize the mechanism of the reconstruction kernel-based batch effect.

## Methods

### Phantom data

The phantom data used in this study, was published by Mahmood et al (Mahmood et al, 2021). To build this phantom, a multimaterial 3D printer (PolyJet Objet 260 Connex 3, Stratasys, Eden Prairie, Minnesota) equipped with Voxel Print software was employed to deposit droplets of UV-curable photopolymer resins in a layer-by-layer manner. Two base photopolymer resins, TangoPlus and VeroWhite, were used, with attenuation ranging from approximately 65 Hounsfield units (HU) to 125 HU at 120 kVp. The printer’s resolution was on the order of 48×84×30 μm, finer than that of typical CT scanners. To replicate contrast differences observed in tumor-specific patterns on patient CT scans, multiple resin droplets were mixed in a single voxel using the Floyd Steinberg dithering algorithm. Four pancreatic ductal adenocarcinoma tumors were printed that captured the different morphological appearances and intensity patterns seen on human CT scans. Prior to dithering, the HU values of each tumor and background were separately converted to double precision intensities within a range of 0 to 1. After 3D printing, the phantom underwent 30 repeat scans using a 64-slice CT scanner (HD750, General Electric, Madison, Wisconsin) with the following parameters: 120 kVp, 0.7 s, 0.984; and the standard, bone, soft, ASIR and ASIRV kernels was applied to the data and two different tube currents were used (100 mAs and 280mAs). (Mahmood et al, 2021)

### Reconstruction kernel effects

Reconstruction kernels are algorithms used during the CT image reconstruction process that affect the appearance of the final image by influencing image sharpness, noise, and the representation of anatomical structures. In this study, multiple convolution kernels were investigated, including standard, bone, soft kernels. These were applied in conjunction with reconstruction algorithms including ASiR (Adaptive Statistical Iterative Reconstruction) and ASiR-V. ASiR reduces the pixel standard deviation by modeling the statistical distribution of noise. ASiR-V includes additional modeling of the system optics and physics to reduce noise while preserving spatial resolution. Each kernel was applied to the phantom scans to observe its effect on the radiomic features. It is crucial to assess the robustness of radiomic features against different reconstruction kernels, as clinical CT scans may employ varying kernels based on specific diagnostic needs. Ensuring that radiomic features remain stable across different kernels enhances their reliability and generalizability in clinical applications (Mahmood et al., 2021).

### Integral tube current

Integral tube current, measured in milliampere-seconds (mAs), is a critical parameter in CT imaging that controls the amount of X-ray photons produced during a scan, directly impacting image noise and radiation dose. In this study, two different integral tube currents, 100 mAs and 280 mAs, were utilized to scan the phantoms. This variation was essential to investigate how changes in integral tube current affect the radiomic features. Evaluating the impact of different tube currents on radiomic features is vital to ensure these features remain consistent despite fluctuations in image noise levels. Robust radiomic features that maintain stability across varying tube currents are more reliable and generalizable in clinical applications (Kalra et al., 2004).

### Radiomic feature extraction

The features were extracted from the original image using the computational environment for radiological research (CERR) (Apte et al., 2018). Standard radiomics features, 282 in total, were extracted from CT images, including first-order statistics (FOS), gray-level co-occurrence matrix (GLCM), gray-level run-length matrix (GLRLM), gray-level size-zone matrix (GLSZM), neighborhood gray-tone difference matrix (NGTDM), and neighboring gray-level dependence matrix (NGLDM). CT image intensities were discretized using a fixed bin width of 5 (HU). The minimum, maximum, mean, mean absolute deviation, and standard deviation for each texture feature were computed over all 13 directions to obtain rotational invariance. A patch-wise volume of 2 × 2 × 2 mm voxels was used for neighborhood gray level dependence features.

### Combat-based batch correction

ComBat was originally developed for genomic data harmonization (Johnson et al, 2007) and more recently has been adapted for radiomics (Fortin et al, 2018; Orlhac et al, 2019). It uses a Bayesian statistical approach to adjust for batch effects while preserving clinically relevant biological variations. The ComBat correction process involves estimating batch effect parameters by modeling data as a combination of batch and biological effects. Specifically, the model can be expressed as (equation 1: eq1):

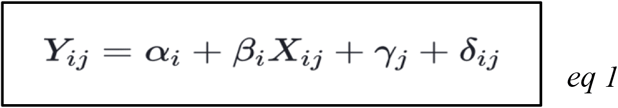

where

*Y*_*ij*_ is the observed feature value for the *j*^*t*h^ sample in the *i*^th^ batch, α_*i*_ *and* β_*i*_ are the parameters that capture the batch-specific effects, *X*_*ij*_ represents the biological variation, *γ*_*j*_ and *δ*_*ij*_ are the batch effect parameters and random error, respectively. These parameters are estimated using an empirical Bayes approach, and the batch effect-adjusted values are obtained as follows:

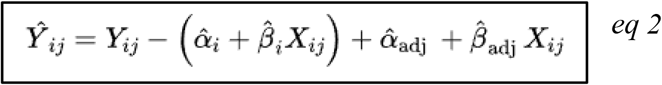

where 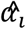 and 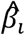 are the estimated batch-specific parameters, 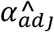 and 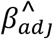 are the adjusted parameters applied to harmonize the data across batches.

### Statistical analysis

#### Assessment of batch effects

Two approaches were employed to evaluate the batch effect. First, principal components analysis (PCA) was performed on the entire set of features as well as on subsets of features from each class. Second, we used visualization methods based on Pearson correlation coefficients to visualize the correlations within each feature class, comparing them across the different kernels and tube currents. This allowed us to assess how the correlations change across different kernels and tube current settings.

#### Robustness of radiomic features

The assessment of feature robustness was based on the concordance correlation coefficient (CCC) of Lin (Lin, 1989). It describes the extent to which paired measurements diverge from perfect agreement, reflecting both systemic differences between repeated measurements and variability. (Lin, 1989; Steichen & Cox, 2002). It’s estmated by the following formula *(eq 3)*:

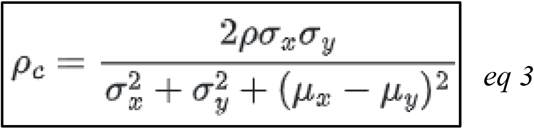

where:

- ρ is the Pearson correlation coefficient between the two variables.
- σ_x_ and σ_y_ are the standard deviations of the two variables.
- μ_x_ and μ_y_ are the means of the two variables.

In this study, to define the robustness, we estimated the CCC between the different kernels and also between the different integral tube current considered. Features was defined to be robust across kernels or integral tube current for CCC ≥ 0.9 as suggested by McBride et al (*McBride et al, 2005*).

#### Software

The data was analyzed using the R software platform (v4.1.3., R core team 2022). We used the yardstick (v1.1.0, Max kun et al) library to estimate the CCC and the *Factoshiny* (v2.5, Vaissie et al) and *corrplot* (v0.92, Taiyun et al) libraries for visualization.

## Results

### Kernel batch effects

#### Kernel batch effects: Visualization using PCA

The visualization of the full features (including all the 282 extracted features) matrix considering the ASIRs, ASIRV, and standard kernel, using the principal components analysis (PCA) on the first two principal components, revealed two clusters *(figure 2)*. The two first components carries almost 63% of the variability in the data matrix. As shown in figure 2, we found a gradient growing from ASIR50 to ASIR10 and from ASIRV5 to ASIRV1 across the first component of the PCA. The PCA on the class specific data matrix, revealed the same trend on GLCM features (*figure 3.c*) and the same trend in the opposite direction where observed in the SZM *(figure 3.d) and* in RLM features *(figure 3.b)*. Even though there appeared some differences in the direction of the gradient across the components, the same trend (from ASIR50 to ASIR10 and from ASIRV5 to ASIRV1) was observed in all the class of features.

**Figure 1:**
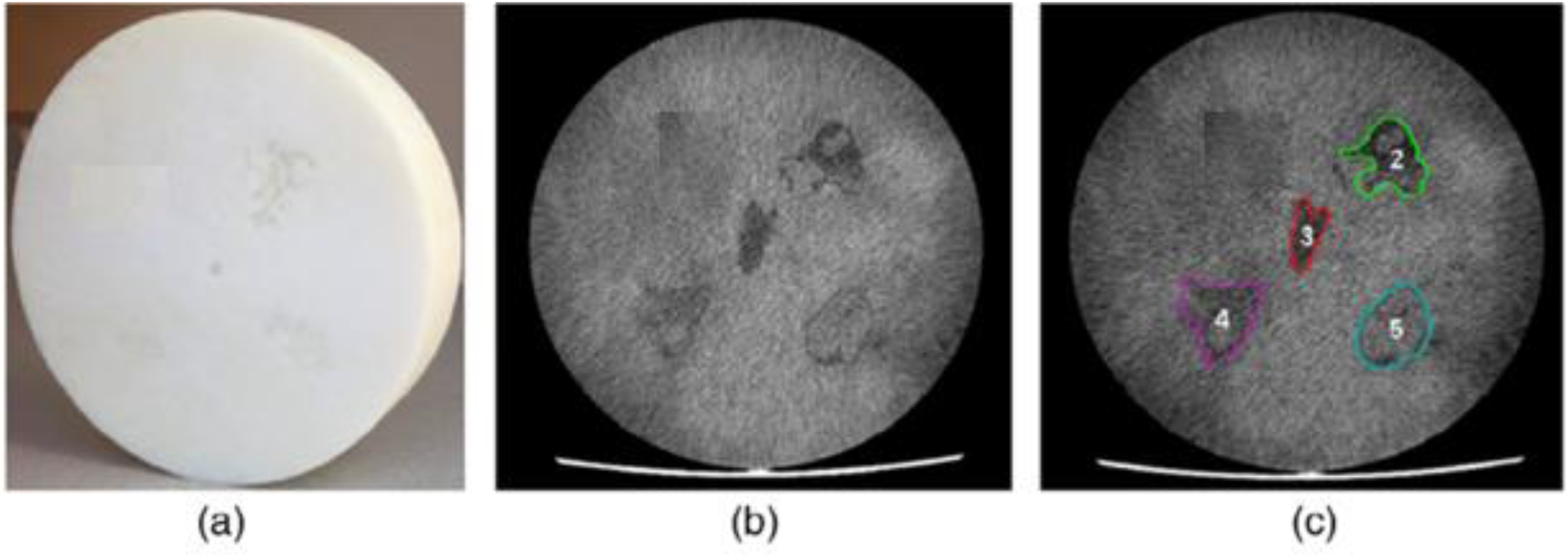
(a) The phantom created using 3D printing. (b) An axial slice produced from a CT scan illustrates the tumors embedded within the surrounding tissue. (c) The outlines of the four pancreatic ductal adenocarcinomas (PDAC). (Mahmood et al., 2021) **Source:** *Published image modified and used with authorization from* (Mahmood et al., 2021)

**Figure 2:**
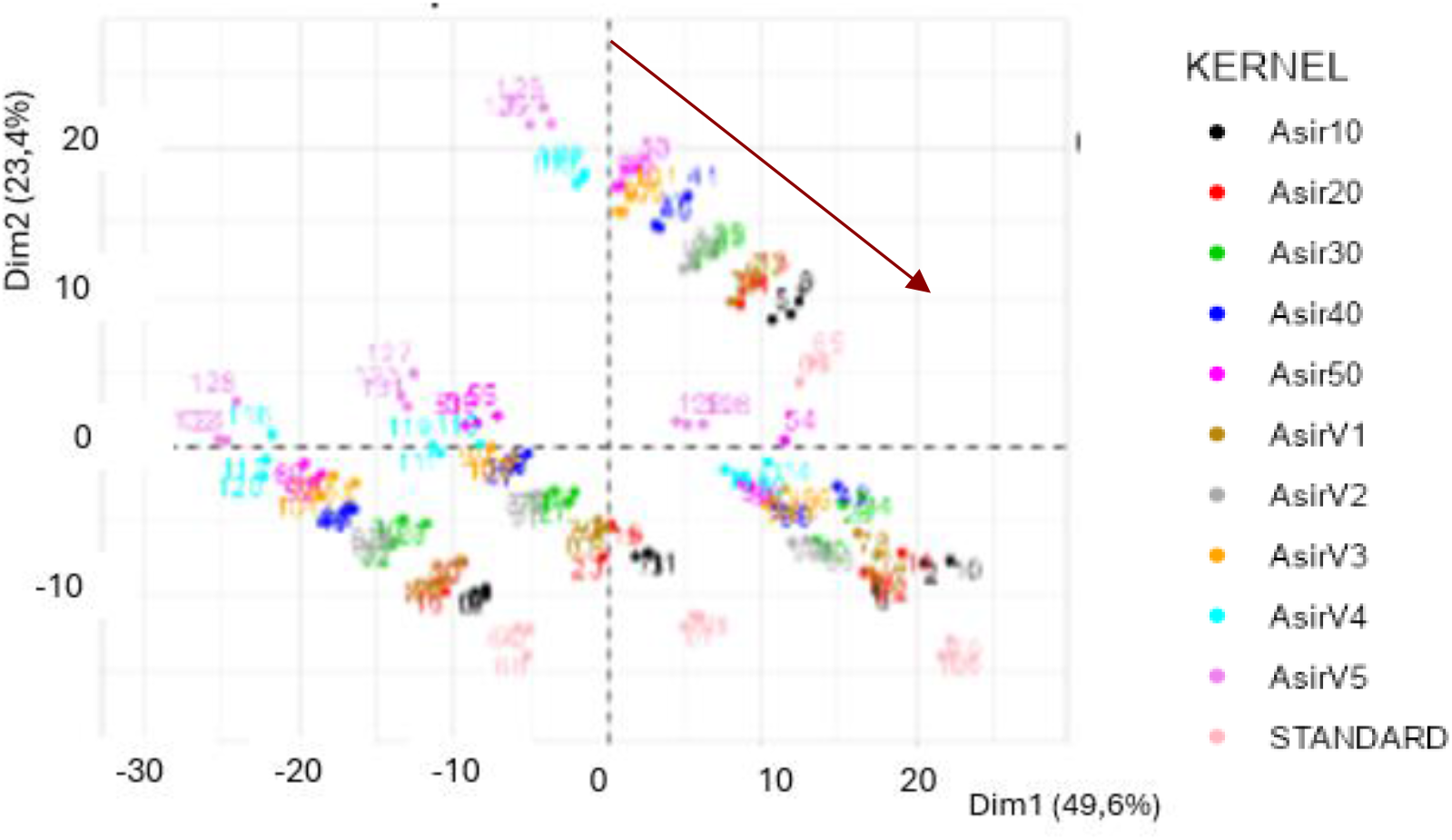
Two first components of the principal component analysis considering for full features matrix colored by kernels ASIRs (10 to 50), ASIRV (1 to 5) and standard kernel. *Numbers are labels of the data points in the matrix. Data extracted from a phantom of Four PDAC identified in the figure by* ***(A)***,***(B)***,***(c)***,***(D)***. *The red arrow represents the trend/ gradient observed between the different kernels*.

**Figure 3:**
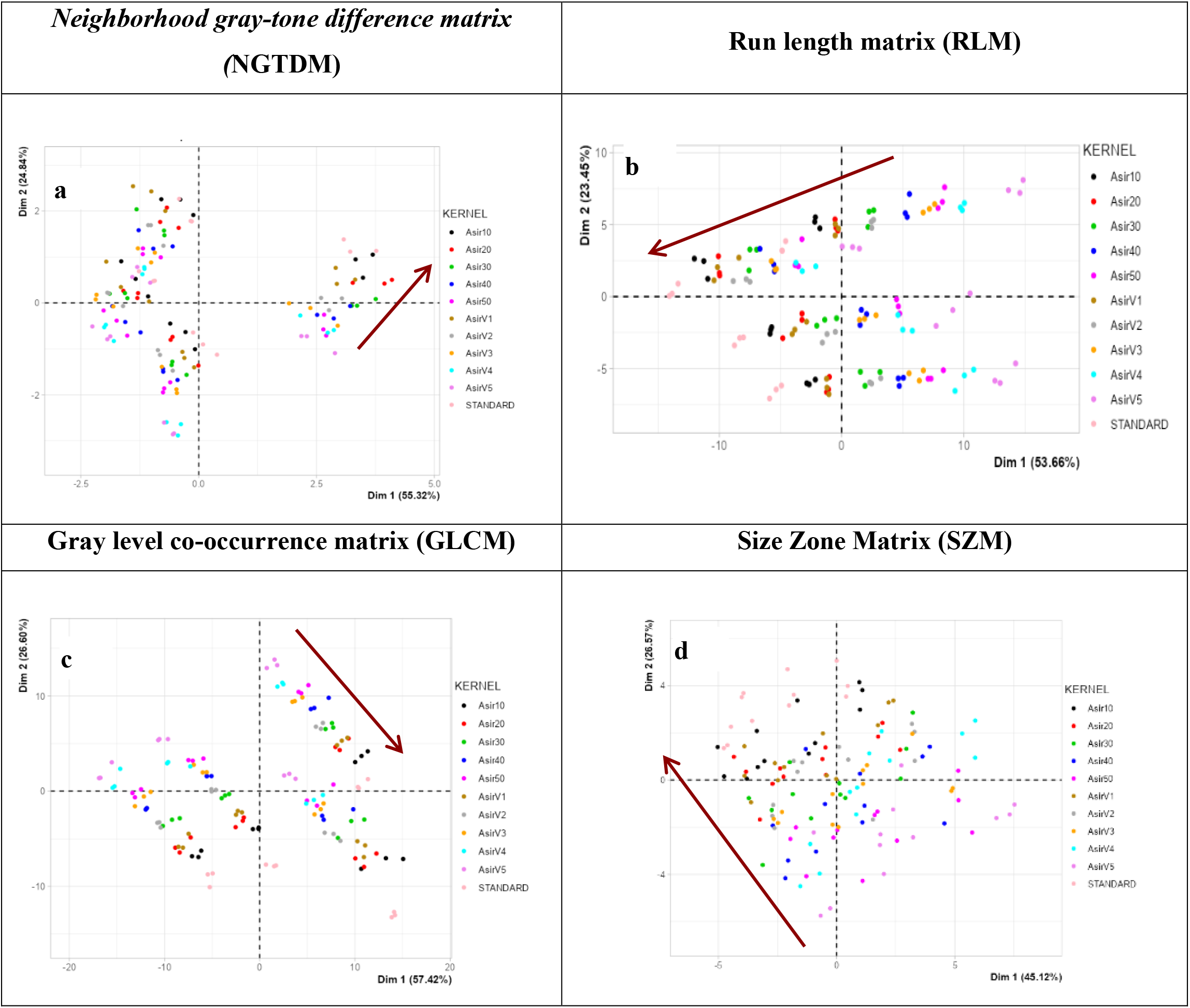
Two first components of the principal component analysis considering for each features class colored by kernels ASIRs (10 to 50), ASIRV (1 to 5) and standard kernel. Data extracted from a phantom of *Four PDAC. (a): represent the results for NGTDM features (17 features), (b): represent the results for RLM features (80 features), (c) represent the results for GLCM features (130 features), (d) represent the results for SZM features (16 features). The red arrow represents the trend/gradient observed between the different kernels*.

#### Kernel batch effects: Correlations between features

As shown in figure 4, concerning the FOS dataset, the feature correlation matrix is still globally conserved with a reduction of the correlation value when comparing the correlation matrix when comparing the results from the Standard kernels to the ASIRs *(figure 4.b)*, ASIRV (*figure 4.d*). In the First order statistics, these changes in the correlation value were most visible in the *skewness* (reduction of the c*orrelation value between skewness and the other features of the First order statistics matrix*).

**Figure 4:**
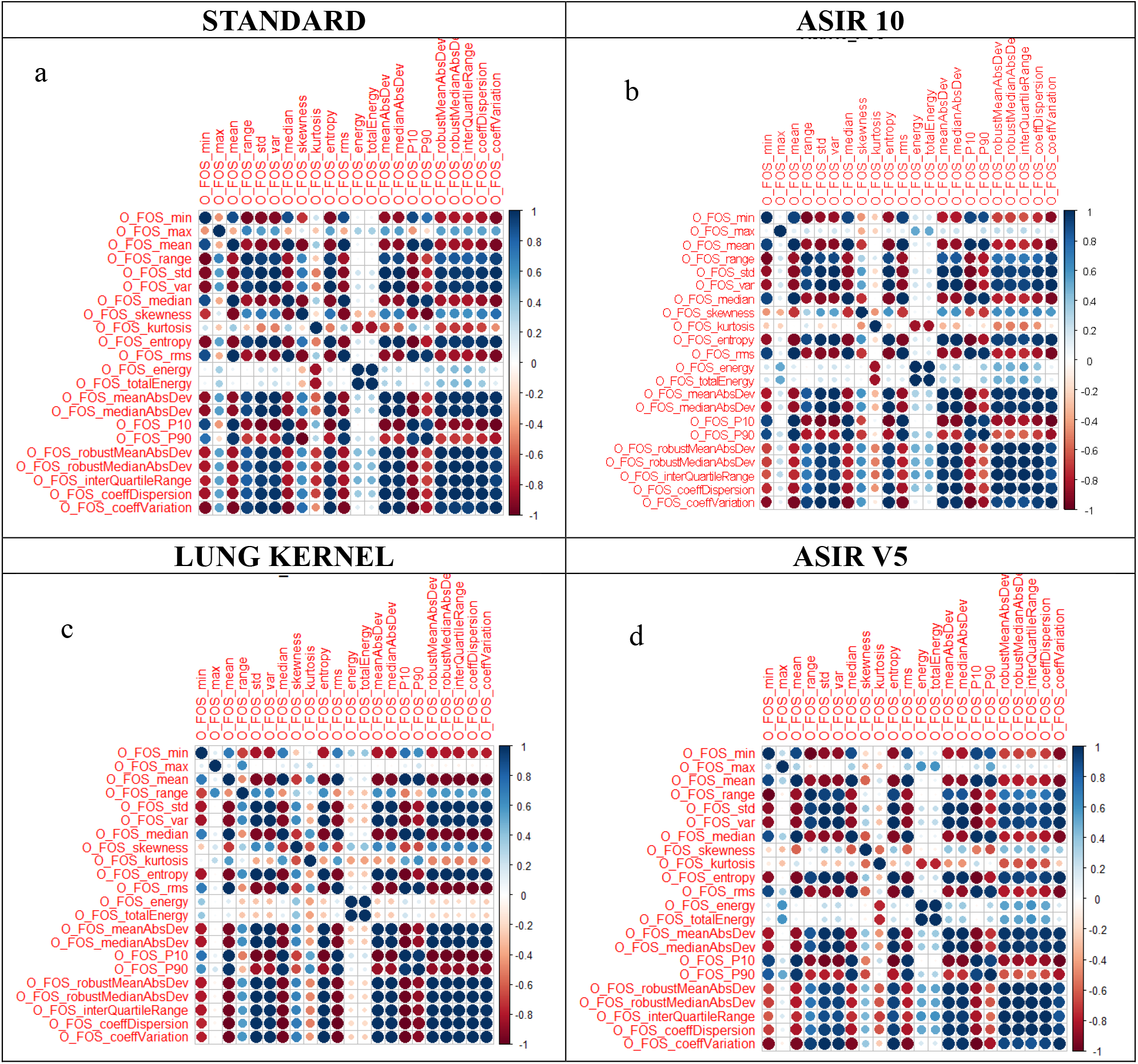
Correlation matrix of the first-order statistics with different kernels (a), standard kernels, (b) ASIR 10, (c) Lung kernel (d) ASIR V5. FOS: First order statistics, O: Original image, P10: percentile 10, P90: percentile 90, var: variance, std: standard deviation, AbsDev: absolute deviation. Min: minimum, max: maximum.

However, important changes in the correlation value appeared in the Bone Kernel and the Lung Kernel *(figure 4.c)*. These changes were most apparent in the correlation value between the *range, kurtosis, skewness* and the other features of the first order statistics. Moreover, the value of the correlation between the energy, total energy, and the dispersion features were improved with the Bone kernel and the Lung Kernel. These findings highlight the ability of the ASIR *(figure 4.b)* and the Standard Kernel *(figure 4.a)*. to conserve the functional relations between the first order statistics.

In the neighborhood gray tone difference matrix, the correlations between the *contrast, the busyness*, the *coarseness and the strength* were unstable with a difference in the sign between different kernels (*Supplementary materials)*.

In addition, in the analysis of the correlation matrix of the NGDLM matrix, notable changes were observed in the *energy, entropy, gray level variance, and gray level non uniform norm*.

### Integral tube current batch effect

#### Visualization using PCA

Analysis of the two tube-currents (100mA Vs 280 mA) considered in this study revealed a global trend in the opposite direction of the first component which carries almost 51.5% of the variability in the data. *(figure 9.a)*. Analyzing the different class of features, the same trend were observed for First order statistics *(figure 5.a)* and and gray level co-occurence matrix *(figure 5.b)*. However in Run length matrix *(figure 5.c)* and size zone matrix *(figure 5.d)* the opposite trends where observed.

**Figure 5:**
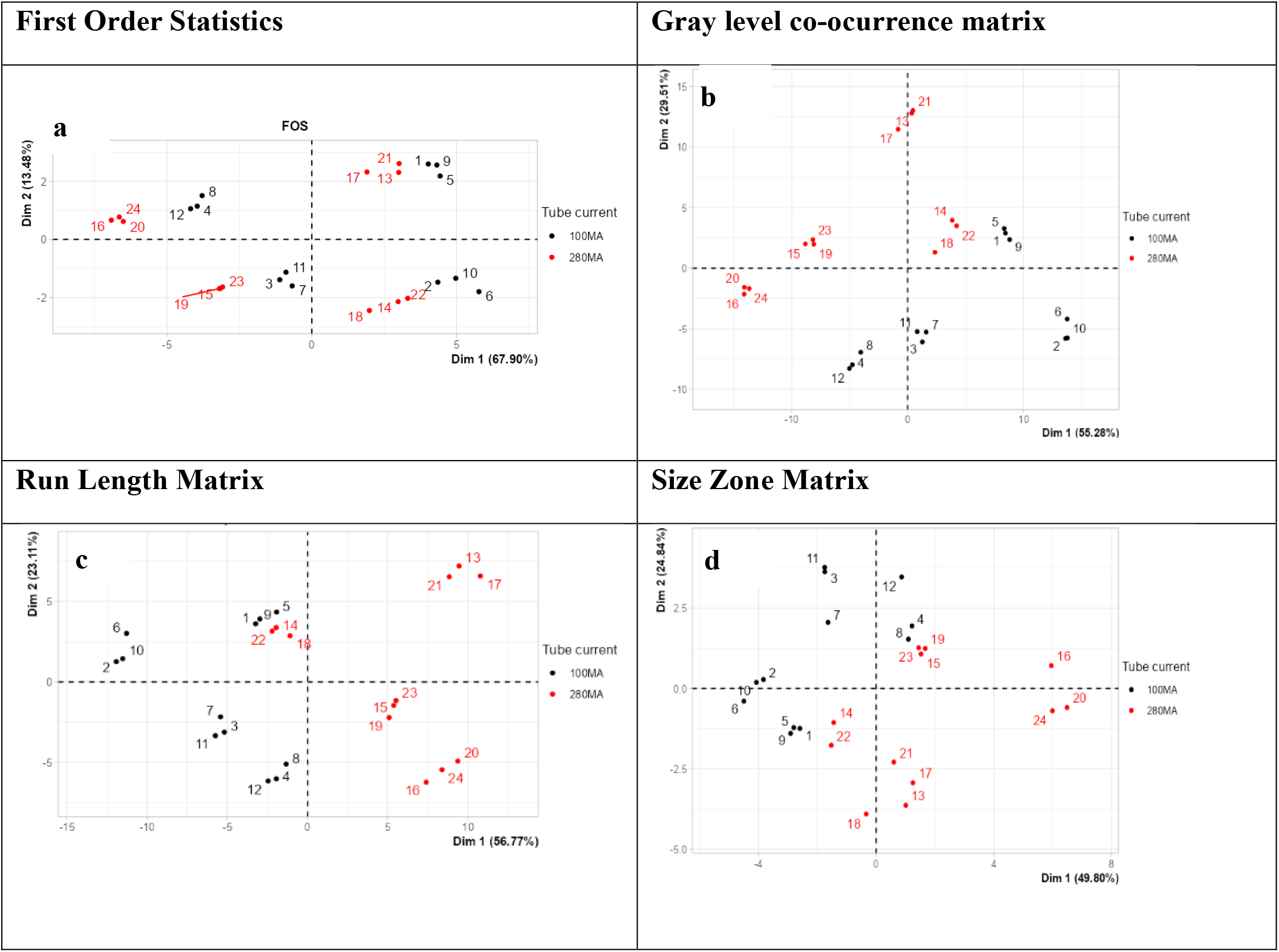
principal component analysis of the different class features colored by the integral tube current. (a) First Order Statistics, (b) Gray level co-occurence, (c) Run length matrix, (d) size zone matrix. Data was extracted from a phantom of four PDAC tumors as described.

#### Correlations between features

As shown in figure 6, the structure of the correlation matrix and the signs of individual correlation values remain largely the same between the 280 mAs and the 100 mAs integral tube current (*figure 6.a vs figure 6.b & figure 6.c vs figure 6.d)*. This highlighted the ability of this type batch effect to conserve the relation between the features of the same class. While the kernel batch effect affected the correlations between the different features, they are not affected by the integral tube current related batch effect.

**Figure 6:**
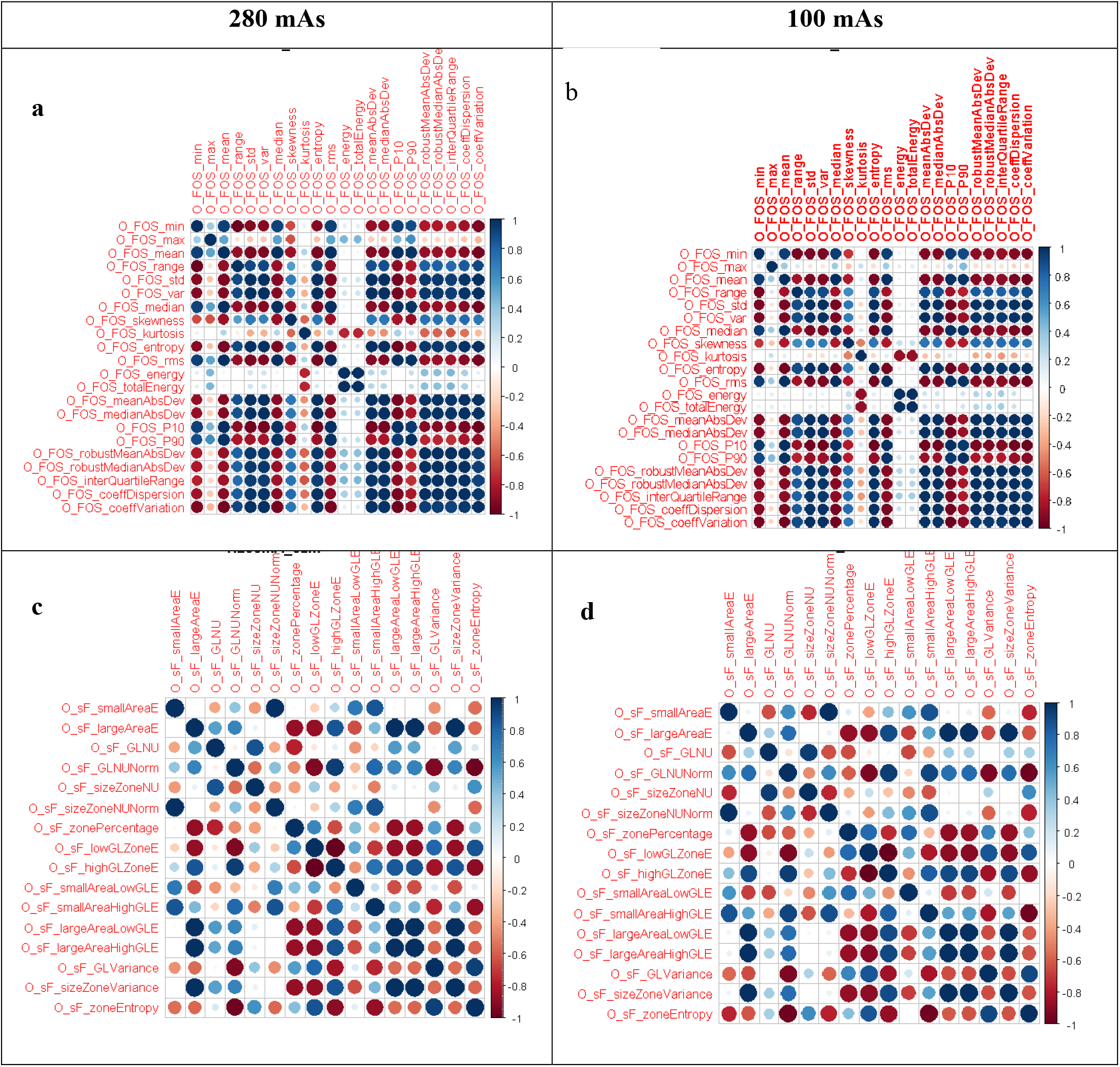
Correlation matrix of the first order statistics with different integral tube current. FOS: First Order Statistics, sF: Size Zone Feature, O: Original image. Data was extracted from a phantom of four PDAC tumors as described. *(a): First Order Statistics features with integral tube current 280 mAs, (b): First Order Statistics features with integral tube current 100 mAs, (c): First Order Statistics features with tube current 280 mAs, (d): First Order Statistics features with integral tube current 100 mAs*.

#### Identification of robust features

Almost 13.6% of the features were robust for all the reconstruction kernels evaluated. As depicted in Figure 7, from ASIR 10 to ASIR 50, there is a decrease in the percentage of robust features between the different ASIR kernels and the Standard kernel. The same trend was observed in the model-based version of the ASIR (*ASIRV*), with a decreasing number of concordances from ASIRV1 to ASIRV5. The first-order statistics were the most robust (22.73%) of all the classes of features. Conversely, no robust features were identified within the SZM and the NGTDM.

**Figure 7:**
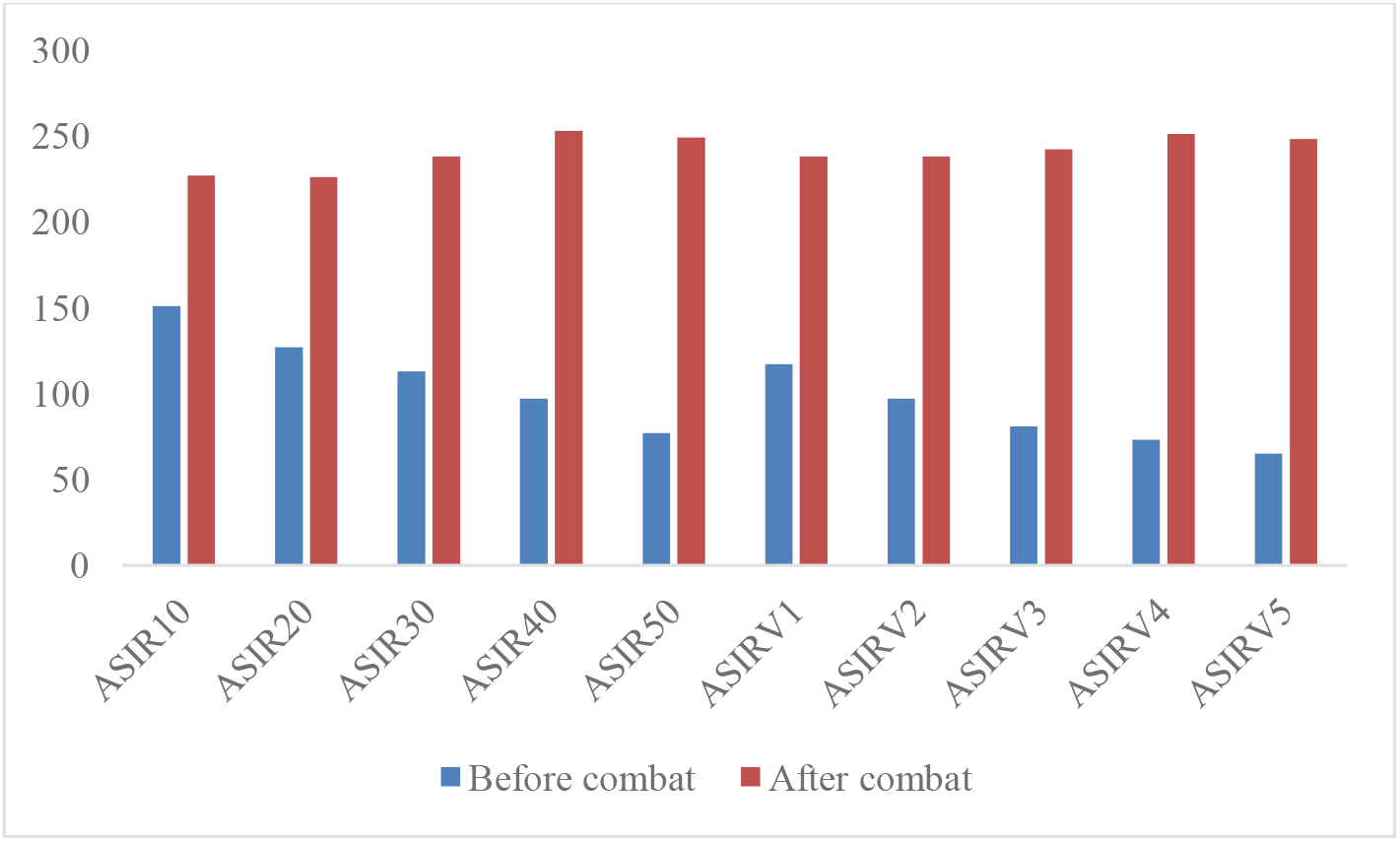
Number of robust features identified for the ASIR (10 to 50) and ASIRV (1 to 5) before and after batch correction. Data was extracted from a phantom of four PDAC tumors as described.

About a quarter (23.7%) of the radiomics features were robust to the integral tube current batch effect. The neighborhood level dependence matrix class was the most robust, with 40% of features classified as robust, while the size zone matrix class had the lowest percentage of robust features (16.25%). Overall, we identified 39 features that were robust to both reconstruction kernel and integral tube current batch effects.

#### Combat corrections for the different batch effects

As shown in Figure 8, the patterns observed in the original dataset *(figure 8.a)* with respect to the kernel disappear after the application of the Combat correction method *(figure 8.b)* on the ASIRs and the standard kernel. However, on the Bone and the Lung kernel, *(figure 8.b)* there was less improvement *(figure 8.b)*. Moreover, as presented in Table 1, the application of Combat to the kernel batch effects significantly improved the number of robust features from *13.7% to 73.2%*. Out of all the features, the first order statistics were still the most robust after correction *(rising from 22.72% to 81.82% robust features)*, while the size zone matrix features and the neighborhood gray level features were the least robust. As presented in figure 9 the application of the combat correction *(figure 9.b)* reduce the distance between the clusters formed by the integral tube current (figure 9.a) even though, there remain a pattern with data points clustered by the integral tube current (figure 9.b). Nevertheless, as shown in Table 2, Combat correction increased the percentage of robust features (*from 23.7% vs 66.55%*).

**Table 1:** Percentage of robust features in the different classes of features before and after kernel batch effects correction *(using Combat-based batch correction)*

| Feature class | Before correction* |  | After combat-based correction |  | Total number of features |
| --- | --- | --- | --- | --- | --- |
|  | Number of robust Features | % | Number of robust Features | % |  |
| SHAPE | 17 | 100.0 | 17 | 100.0 | 17 |
| FOS | 5 | 22.7 | 18 | 81.8 | 22 |
| GLCM | 9 | 6.9 | 95 | 73.1 | 130 |
| RLM | 6 | 7.5 | 59 | 73.7 | 80 |
| NLDM | 0 | 0.0 | 2 | 40.0 | 5 |
| NGTDM | 2 | 11.8 | 12 | 70.6 | 17 |
| SZM | 0 | 0.0 | 7 | 43.7 | 16 |
| SHAPE | 39 | 13.6 | 210 | 73.2 | 287 |

\* Data extracted from a phantom of Four PDAC. ASIRs, ASIRV and standard kernels were applied to the phantoms. ‘Robustness’ is defined in the methods section.

**Table 2:** Percentage of robust features in the different classes of features before and after integral tube current batch effects correction *(using Combat-based batch correction) with standard kernel*.

| Feature class | Before Correction* |  | After COMBAT-based correction |  | Total number of feature |
| --- | --- | --- | --- | --- | --- |
|  | Number of robust Features | % | Number of robust Features | % |  |
| SHAPE | 17 | 100.0 | 17 | 100.0 | 17 |
| FOS | 7 | 31.8 | 18 | 81.8 | 22 |
| GLCM | 24 | 18.5 | 87 | 66.9 | 130 |
| RLM | 13 | 16.2 | 50 | 62.5 | 80 |
| NLDM | 2 | 40.0 | 3 | 60.0 | 5 |
| NGTDM | 3 | 17.6 | 10 | 58.8 | 17 |
| SZM | 2 | 12.5 | 6 | 37.5 | 16 |
| Total | 68 | 23.7 | 191 | 66.5 | 287 |

\*Data extracted from a phantom of Four PDAC. Two tube currents 100mAs and 280 mAs with standard kernels were applied to the phantoms. Robust features a meant to be robust across the different integral tube current as defined in the methods section.

**Figure 8:**
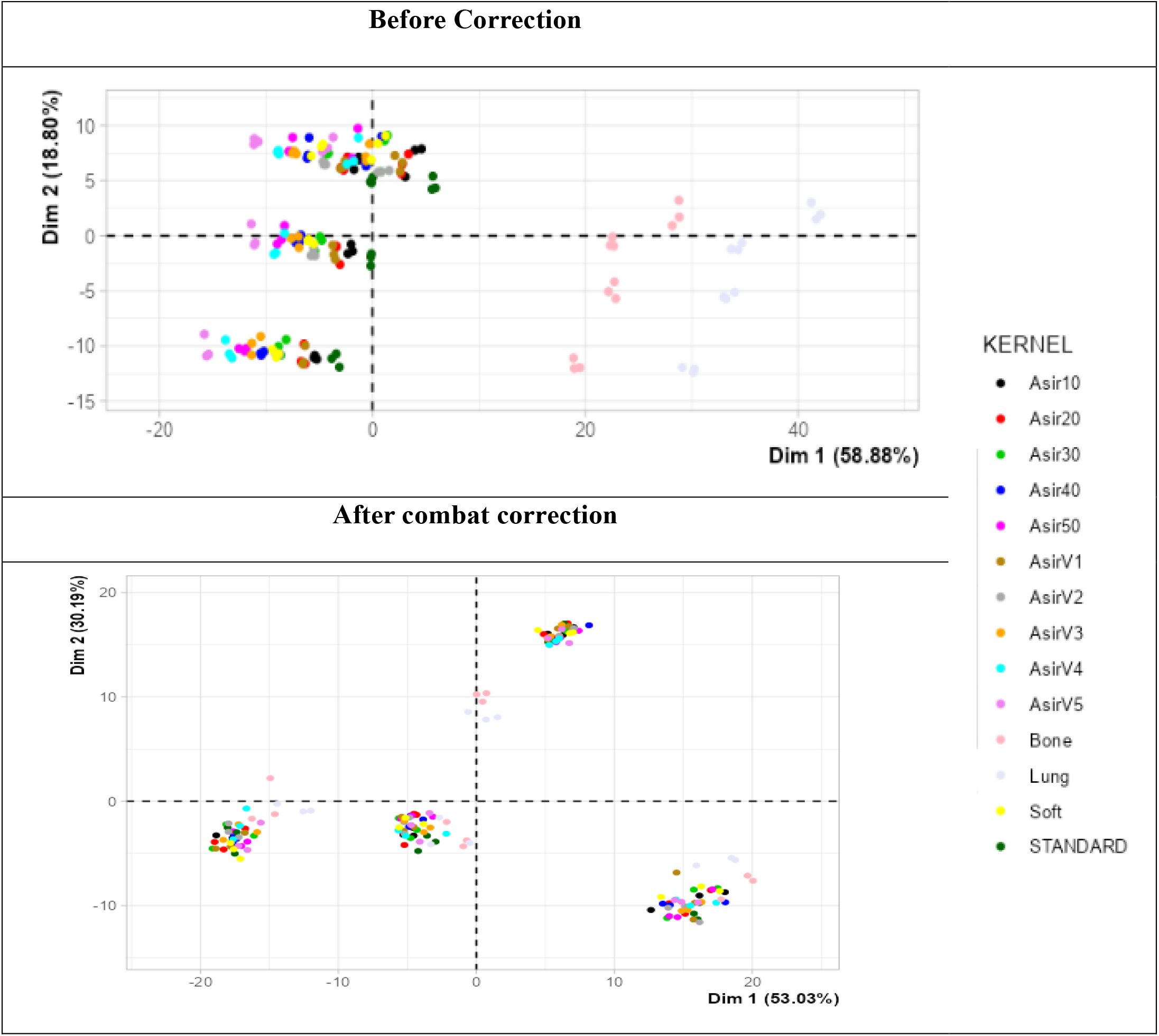
Two first components of the PCA applied to the full features matrix colored by reconstruction kernels (a) Before correction and (b) After combat based data correction. *Data extracted from a phantom of Four PDAC*.

**Figure 9:**
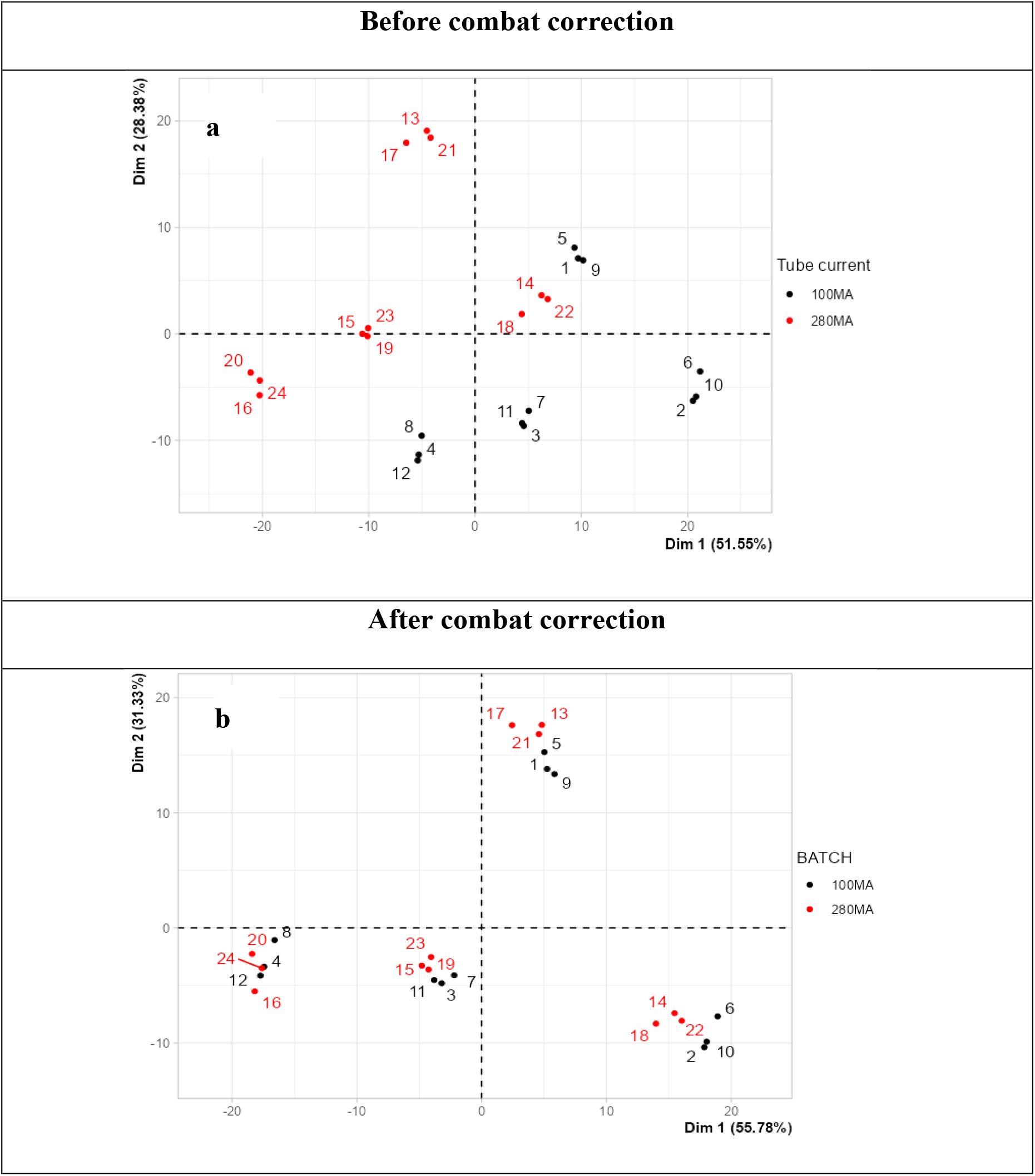
Two first components of the PCA applied to all the features with different tube currents (100 mAs vs 280 mAs) (a) before and (b) after Combat batch effects correction colored by integral tube current. *Data extracted from a phantom of Four PDAC*.

## Discussion

Previous investigators have shown that some features are robust to integral tube current and possibly other reconstruction parameters, including reconstruction kernel choice. Using a tumor-imprinted phantom allowed us to obtain and analyze a dataset where reconstruction kernel and tube current effects and reproducibility could be studied in detail. In this study, we have identified 39 features robust to both the reconstruction kernel and the integral tube current batch effect. The choice of kernel affects the concordance and the correlation matrix between features of the same class, while the integral tube current does not significantly affect the correlation matrix of features in the same class. The application of Combat batch correction improves the proportion of robust features for each of the considered batch effects. Visual inspection of the principal components reveals that Combat’s performance is better when applied to kernel batch effects from the ASIR, ASIRV, and standard kernels, while Bone and Lung kernel data harmonization still needs improvement. When correcting for integral tube current batch effects, Combat improves data harmonization by a modest degree, although clustering/batch effects can still be observed.

The ability of Combat to improve the robustness of features has already been demonstrated in various studies (Eshaghzadeh Torbati et al., 2021; Horng et al., 2022; Mahon et al., 2020). However, its performance is subject to limitations related to the structure of radiomics data. In our study, we found that combat harmonization performs differently depending on the type of batch effect considered. This can be explained by differences in the functional forms and the mechanisms behind those batch effects, which have not been explored well in the literature.

Analysis of the robust features before correction revealed that the central-tendency parameters in the first-order statistics were robust to kernel and integral tube current batch effects. This robustness suggests that these batch effects do not significantly influence the central tendency and the global dispersion of the voxel intensity, while by contrast, the dispersion parameters and skewness were changing. The latter two measures detect variations in the shape of the distribution, suggesting changes in dispersion and symmetry of the voxel intensity distribution between batches. Moreover, this contrast between the central tendency parameters and the dispersion and shape parameters highlights a non-linear batch effect, with specific influence depending on image region. Those regions change according to the kernels considered. Within the the gray level co-occurrence matrix feature class, the maximum, mean, minimum and standard deviation of the “high gray level run emphasis”, “low gray level run emphasis” and “run lengths non uniformity” were robust. The consistency of this measure indicates that the variation in run lengths between batches remains relatively constant. More broadly, the stability of these parameters suggests that the kernel and the integral tube current batch effects do not significantly affect the global distribution of gray levels, the prevalence of high or low gray level runs, and the irregularity of run lengths while the dispersion parameters are changing.

While there exist multiple data driven and deep learning-based corrections methods, (Fortin et al, 2018; Homg et al 2022, Orlhac et al,) a critical remaining issue is that the correction itself is not transparent. As demonstrated with ComBat, these batch effect correction methods can work for some types of batch effects, but not other types. Given that multiple types of batch effects are found in radiomics, the fact that correction methods can account for some types and not for other types of batch effects is a continuing obstacle to reliance on radiomic or deep learning models (Li et al, 2021; Van velden et al 2016; Zhao et al, 2014). Most of the harmonization methods introduced thus far apply some global correction technique, while this study revealed that the kernel and integral tube current batch effects conserve the global forms and instead affect specific regions in the data through a nonlinear and non-constant functional transformation. In this context, the application of a linear correction may overcorrect some features and lead to bias in the results.

Although a radiomic phantom makes comparable scanning possible, it does not completely substitute for studies in human datasets. Another limitation of this study is that data were extracted from one scanner alone. Hence, over-interpretations should be avoided. Future directions should include validating these findings in patient datasets and further exploring the non-robust and robust features to extract the functional forms of different types of batch effects. Also, given the lost context once features are derived, methods should be explored to improve harmonization by operating at least partly in the image space.

## Conclusions

About 14% of the 282 investigated standard radiomic features were robust to kernel and integral tube current batch effects without any batch correction. Combat improved this to 70%. However, the variability of many radiomic features due to kernel reconstruction choices was not improved adequately by ComBat, particularly bone and lung kernel results. integral tube current differences, in contrast, were better corrected by Combat. Further work is required to determine if increased harmonization is possible for a larger range of radiomic features, possibly applying image based methods.

## Acknowledgements

We acknowledge the late Prof. Allen Tannenbaum for initiating this collaboration. The stay of Elfried Salanon was supported by the I-SITE project & PFEM/INRAE, and his PhD project is fully supported by the University of Geneva and the Digit-Bio metaprogram of INRAE. This research was also supported by the Breast Cancer Research Foundation and the Simons Foundation.

